# A mathematical model exhibiting the effect of DNA methylation on the stability boundary in cell-fate networks

**DOI:** 10.1101/2019.12.19.883280

**Authors:** Tianchi Chen, M. Ali Al-Radhawi, Eduardo D. Sontag

**Author notes:** These authors contributed equally (Tianchi Chen, M. Ali Al-Radhawi).

## Abstract

Cell-fate networks are traditionally studied within the framework of gene regulatory networks. This paradigm considers only interactions of genes through expressed transcription factors and does not incorporate chromatin modification processes. This paper introduces a mathematical model that seamlessly combines gene regulatory networks and DNA methylation, with the goal of quantitatively characterizing the contribution of epigenetic regulation to gene silencing. The “Basin of Attraction percentage” is introduced as a metric to quantify gene silencing abilities. As a case study, a computational and theoretical analysis is carried out for a model of the pluripotent stem cell circuit as well as a simplified self-activating gene model. The results confirm that the methodology quantitatively captures the key role that methylation plays in enhancing the stability of the silenced gene state.

## 1 Introduction

Cell-fate determination in developmental biology is a multi-step biological process in which cellular functions are modified or specialized, ultimately resulting in a differentiated or proliferative state. A central role in this process is played by the dynamics of a core network of genes, often referred to as a *cell-fate network* (CFN). The expression levels of these genes can trigger a cascade of events that determine, in principle irreversibly, the fate of a given cell along a specific lineage. The metaphor of marbles rolling down a hill whose shape is shaped by a CFN (the *Waddington landscape* [51, 54]) provides a way to visualize this process. Well-known examples of CFNs are the PU.1/GATA.1 gene regulatory network in hematopoietic progenitors [10, 19], the pluripotent stem cell network [19, 11], and a circuit including SNAIL, miR-34, ZEB1, and miR-200 that regulates epithelial-mesenchymal transitions in tumor metastasis [37].

Phenotypes associated to CFNs had classically been seen as irreversible. However, in their pioneering work [47], Takahashi and Yamanaka, artificially induced a pluripotent state in mouse somatic cells through a process of overexpression of Oct3/4, Sox2, c-Myc, and Klf4. This success notwithstanding, simply overexpressing genes has been experimentally found to be grossly inefficient [40]. Hence, there has been a great interest in obtaining a quantitative theoretical understanding of CFNs in order to guide the process of reprogramming and increase its efficiency [34, 15]. Such a quantitative understanding could have a huge impact in the field of regenerative medicine and stem cell therapy [35, 53].

Traditionally, CFNs have been modeled within the wider theoretical framework of Gene Regulatory Networks (GRNs) [25, 28]. A GRN is defined as a set of genes, each expressing a protein. The expressed proteins can act as Transcription Factors (TFs) by binding to the various promoters in the network to inhibit or enhance expression of the corresponding genes [3, 14]. When used to model a CFN, a GRN must be able to display *multistability*. This means that the asymptotic expression levels of genes can settle on multiple distinct steady states, each corresponding to a distinct cell lineage. A well-known multistable GRN is the *toggle switch* [22] whose architecture consists of two mutually inhibiting genes. Similar architectures can also give rise to multistability and have been subject of great interest [54].

Despite their versatility and variety of applications, GRN models of CFNs do not typically account for epigenetic effects, such as DNA methylation, histone modifications, or chromatin remodelling. There has been both theoretical [30, 20, 21] and experimental [8] works aiming at the understanding of how epigenetic regulation and gene regulation interact. It is well-known that CFNs employ epigenetic regulation as a mechanism to ensure the irreversibility of the cell lineage [48]. DNA methylation, in particular, has been well-studied in the context of developmental CFNs [32]. Methylation is a highly heritable, hard to reverse, and robust silencer of genes [6].

In this work, we develop mathematical models of GRNs that incorporate DNA methylation, and thus can quantitatively explain its effect on gene silencing. As a metric to quantify the gene silencing abilities of DNA methylation, we consider the shift in the stability boundary of the basin of attraction (BoA) of the silenced steady state. More precisely, we define the BoA percentage (BoAp) of a steady state as the volume fraction of a predetermined region of the state space that is occupied by the BoA. Equipped with this metric, one can then computationally and theoretically study the sensitivity of the BoAp to parameters such as methylation rates or the time scale of methylation.

We carry out this program computationally, illustrating it with a three-gene model of the pluripotent stem cell network, and quantifying how methylation effectively increases the BoAp compared to a standard GRN. In addition, and in order to gain understanding, we also consider an ideal single self-activating gene, for which a more theoretical study is possible. We view our work as a first step towards integrating epigenetic mechanisms into the standard GRN paradigm studied in systems biology.

### Basic biological concepts and definitions

#### DNA methylation as epigenetic regulation

DNA methylation is one of the main epigenetic regulation mechanisms studied in modern molecular cell biology. Methylation plays a crucial role [13] in understanding the dynamics of gene silencing. It is associated with methylation of Cytosine-Phosphate-Guanine (CpG) islands, which are regions of DNA with a high G+C content and a high frequency of CpG dinucleotides relative to the bulk genome [39]. Methylation of promoter regions directly affects TF-promoter binding kinetics, and is often associated with transcriptional repression [42].

The current understanding of the DNA methylation cycle is shown in Figure 1. As depicted, cytosines at DNA promoter regions are initially methylated by DNA methyltransferase (DNMT) to become 5-methylcytosine (5mC). The methylated gene promoter is further oxidized to 5-Hydroxy Methylcytosine (5hmC). The oxidized form of the cytosine can further be oxidized by the TET protein into the 5fC and 5caC forms. The cycle closes by a TET/TDG (thymineDNAglycosylase)/BER (base excision repair)-dependent pathway that restores the unmodified cytosine state [31].

**Figure 1:**
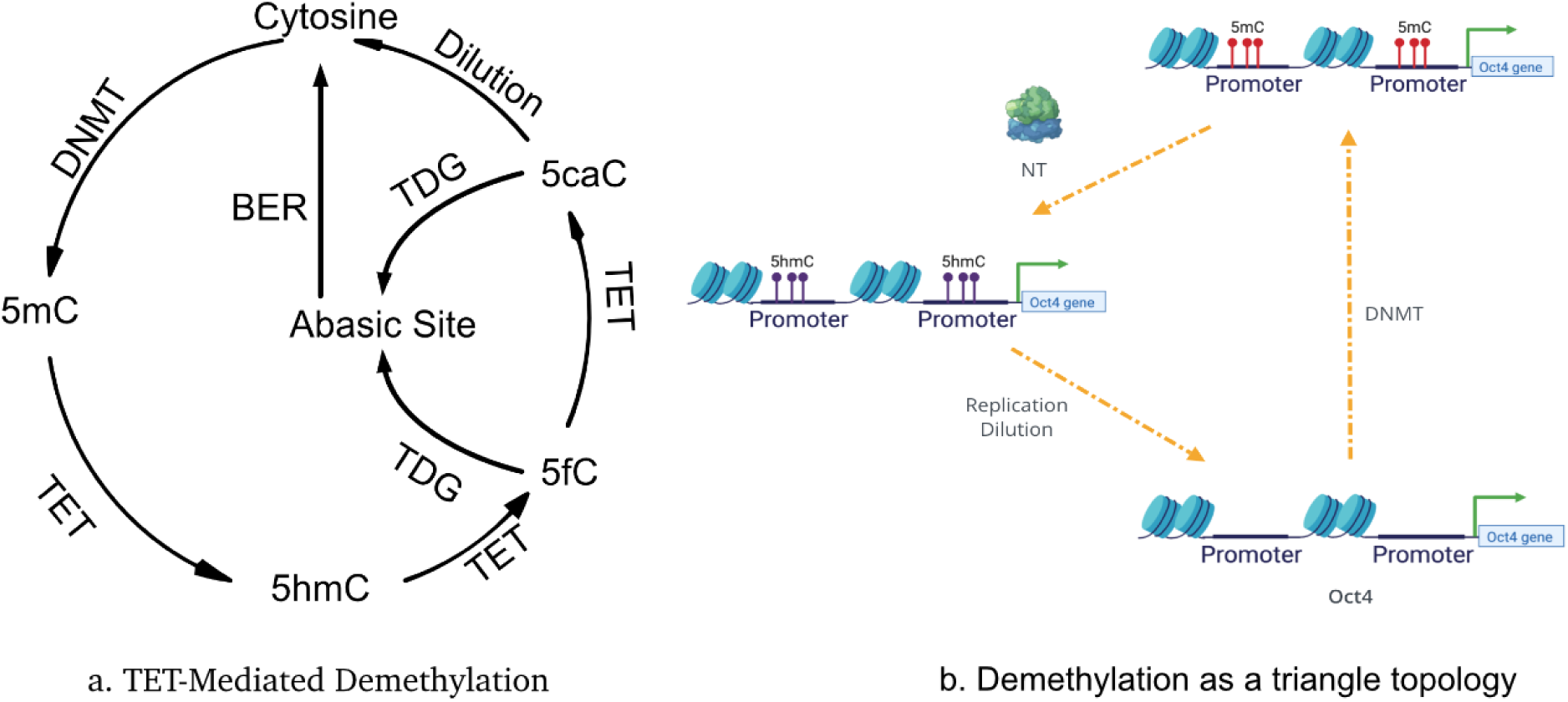
The Methylation Cycle. (a) The TET-mediated Cytosine Demethylation Cycle. (b) A simplified de-methylation cycle for Oct4 in the PSCC.

Experiments have shown that hydroxymethylation and formylation are relatively stable cytosine modifications in genomic DNA of both dividing and nondividing cells [5, 4]. The 5hmC and 5fC forms in this enzymatic oxidation process are observed to be comparatively longer lived, though still transient, states. Moreover, it also has been shown [31] that the pool size of the transient oxidized cytosine 5hmC is significantly larger than the other forms of oxidized cytosines (5fC, 5caC). Therefore, in this work, we further simplify the DNA demethylation cycle to the cycle in Figure 1-b which consists of just three major steps, which we will refer to as the “**triangle topology**.” The cycle starts with an initial de novo methylation on the unbounded gene promoter, which is followed by a TET oxidization enzymatic process oxidizing 5mC to 5hmC. The last stage corresponds to a replication-dependent dilution bringing 5hmC back to the unbound promoter state.

The underlying DNA methylation dynamics is crucial for understanding how epigenetics regulates CFNs. DNA methylation plays a central role not only in early embryogenesis, but also in maintaining the correct pattern of methylation along DNA in somatic cells. As a model system of great interest, we review next how the pluripotent stem cell circuit (PSCC) is affected by the DNA methylation cycle.

#### The Pluripotent Stem Cell Circuit (PSCC)

Some of the key Transcription Factors (TFs) that are identified as being crucial to the PSCC include Nanog, Oct4, Sox2, Klf4, and the TET protein family [47, 31]. Many of such TFs are “pioneering” transcription factors that are able to bind directly to the condensed chromatin. According to the induction experiments in [47], the overexpression of the aforementioned TFs can lead to the induction of the pluripotent state. For the PSCC, the role of each TF in the GRN has been extensively studied [9], and much research has been devoted to elucidating the interplay between the Nanog, Oct4 and TeT1 proteins [50, 16]. Recent experiments regarding the induction of the pluripotent state in PSCCs have highly heterogeneous results [49, 40]. There is evidence [50] that Oct4 over-expression is sufficient to induce the pluripotent state from somatic cells. More recent research [12, 43] has shown that, at a mechanistic level, Nanog is able to guide TET protein to bind to a particular region of the methylated region on the chromatin via the formation of a Nang-TET compound that is actively involved in the DNA demethylation process.

Notably, DNA methylation of the Oct4 gene plays a key role in the regulation of the PSCC. In this work, we model the interplay between epigenetic regulation via the Nanong-guided TET-mediated active demethylation cycle and the role of pioneering transcription factors such as Oct4 and Nanog. A mechanistic view of this interplay helps one understand how emergent cell states can arise phenotypically from the underling genotypic level interaction. Although the full modeling of epigenetic regulation at the molecular level would involve many additional epigenetic regulation mechanisms beyond methylation, such as histone modifications [31], the timescale of DNA methylation is relatively slow [44], and methylation plays a major role in the coarse grained picture of the process of the induction of the pluripotent state which we are mostly interested in.

#### Basin of Attraction percentage (BoAp)

Using high dimensional attractors, and specifically stable steady states, to represent cell fates in the epigenetic landscape is a widely used paradigm for studying cell fate [28, 33]. To further quantify the stability boundary of such attractors, researchers have been using the concept of basin of attraction which is standard in dynamical systems when analyzing multi-stable systems, and has recently been systematically used by the physics community in that context [36]. In our simulation analysis, we adopt this concept to quantify the stability boundary of the somatic cell state. In a bistable one-dimensional model, the BoAp of the smaller steady state *s*_0_ can be defined as the distance between *s*_0_ and the unstable state relative to the length of a segment of interest. However, this is not an accurate measure for high dimensional systems. Therefore, we adopt a *volumetric* definition for the BoA. We start by fixing a region of interest in the state space. With respect to such a region, BoAp can be defined as follows:

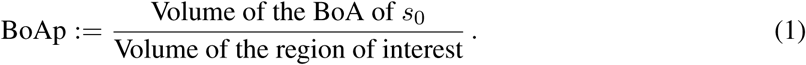

Equipped with this definition, we can study the effect of methylation and demethylation constants on BoAp. Our subsequent results will show that the BoAp of the silenced steady state increases as the methylation rate increases, and decreases as demethylation rate increases.

## 2 Results

### 2.1 The PSCC with epigenetic regulation

In our full model description of the PSCC, the network can be viewed as made up of two parts. The first part is the gene-TF regulatory network [7] which is a core underlying network defining how TFs bind and unbind to DNA as shown in Figure 2 without the shaded area. The main regulatory sub-network for the activation of the Oct4 and Nanong genes has the diamond topology as depicted in Figure 3. Both the Oct4 and the TET gene promoter binding mechanisms will follow the diamond topology, which has two pathways that lead to the activation of the gene. Oct4 protein self-binds to the first site while the Nanog-guided TET protein complex binds to the second site. The promoter is active only when both sites are occupied. In this GRN, the Nanog gene is activated by the Oct4 gene. Furthermore, Nanog and TET form a heterodimer which models TET guiding Nanog to the target promoters.

**Figure 2:**
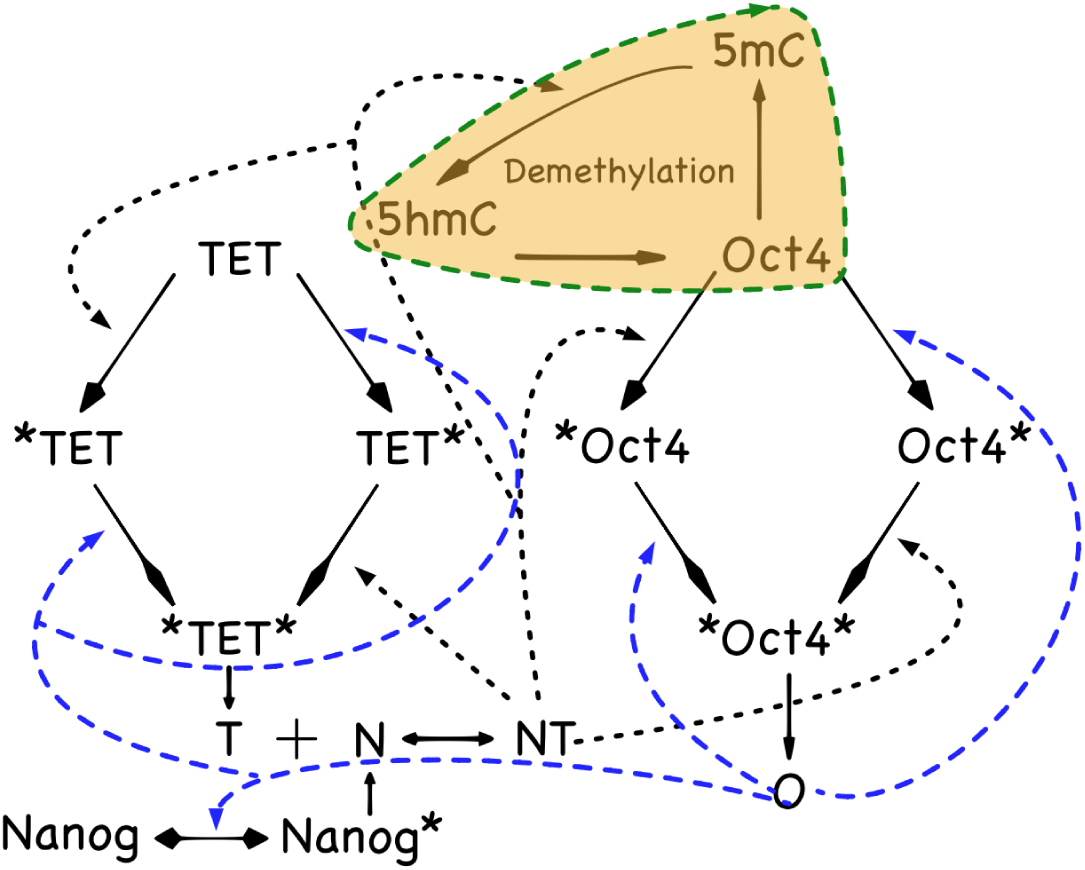
A detailed diagram of the PSCC network showing N, T, O interactions. TET, Oct4, Nanog denote the unbound genes. ^*^TET and ^*^Oct4 denote NT binding to the respective promoter sites, TET^*^ and Oct4^*^ denote Oct4 bound to the respective promoter sites, and ^*^TET^*, *^Oct4^*^ denote both NT and Oct4 binding to the respective promoter sites. The black dashed line indicates binding locations of the Nanog-guided TET protein complex NT, while the blue dashed line indicates binding locations of Oct4. The colored shaded area emphasizes the DNA demethylation cycle

**Figure 3:**
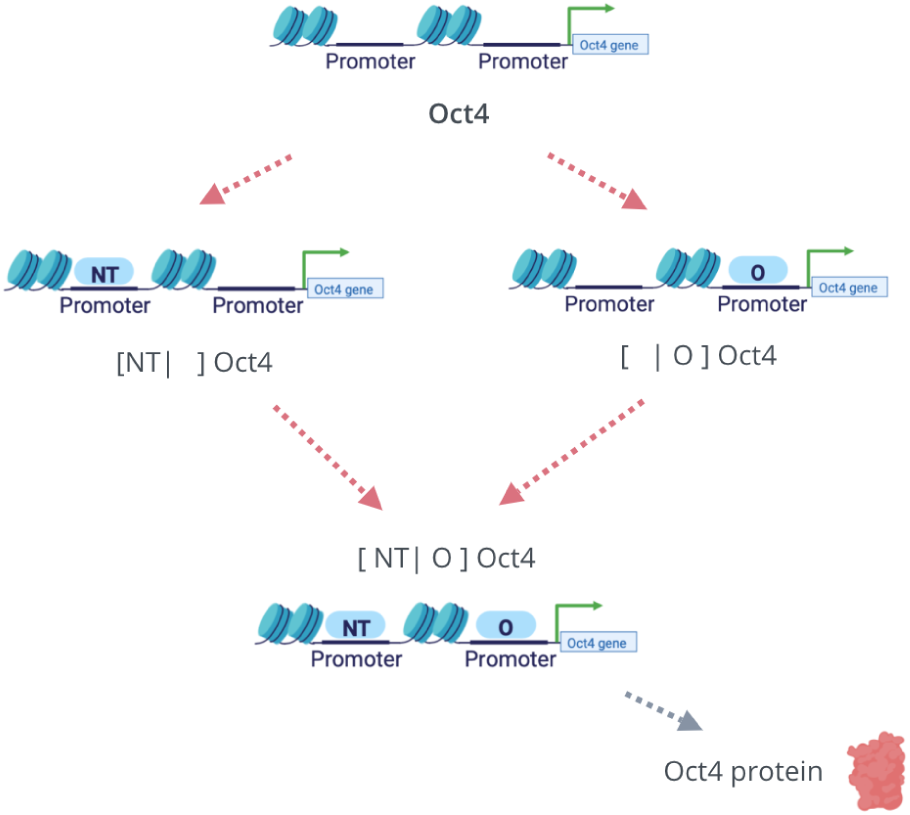
The **Diamond Topology** for double site TF-gene binding. In our model, it describes both Oct4 and TET promoters binding mechanism. Considering Oct4 as an example, there are two pathways for the activation of its promoter. Each of NT and Oct4 can bind to the corresponding Oct4 binding site. When both NT and Oct4 bind to the promoter region of the Oct4 gene, the Oct4 promoter will be activated.

The GRN with DNA demethylation is summarized in Figure 2. The DNA demethylation cycle with the triangle topology can be further simplified into two states, which are shown in Figure 4. The parameter *γ* is thought of as the effective methylation rate and the parameter *θ* as the effective demethylation rate. The CRN for the PSCC with a two-state methylation cycle can be written as in Table 1. (For simplicity, we omit explicit variables for mRNA and other intermediate steps such as protein maturation and post-translation modifications.) We model this network deterministically through a system of ODEs. The variables in this system can be thought of as mean numbers for the various species in a population.

**Table 1:** The CRN model of the PSCC. Reactions have been grouped into eight modules and labeled from R1 to R18

|  |  |  |  |
| --- | --- | --- | --- |
| Oct4 promoter<br>(R1-R4) | $D_{00}^O + [NT] \xrightleftharpoons[K_{nt} * a_{nt}]{a_{nt}} D_{10}^O$ $D_{01}^O + [NT] \xrightleftharpoons[K_{nt} * a_{nt}]{a_{nt}} D_{11}^O$ $D_{00}^O + O \xrightleftharpoons[K_O * a_O]{a_O} D_{01}^O$ $D_{10}^O + O \xrightleftharpoons[K_O * a_O]{a_O} D_{11}^O$ | TET promoter<br>(R5-R8) | $D_{00}^T + [NT] \xrightleftharpoons[K_{nt} * a_{nt}]{a_{nt}} D_{10}^T$ $D_{01}^T + [NT] \xrightleftharpoons[K_{nt} * a_{nt}]{a_{nt}} D_{11}^T$ $D_{00}^T + O \xrightleftharpoons[K_O * a_O]{a_O} D_{01}^T$ $D_{10}^T + O \xrightleftharpoons[K_O * a_O]{a_O} D_{11}^T$ |
| Nanog promoter<br>(R9) | $D_0^N + O \xrightleftharpoons[K_O * a_O]{a_O} D_1^N$ | Nanog-TET<br>Dimer (R10) | $N + T \xrightleftharpoons[K_d * a]{a} [NT]$ |
| Protein decay<br>and production<br>(R11-R13) | $N \xrightarrow{\delta} \emptyset; D_1^N \xrightarrow{\alpha_N} D_1^N + N$ $T \xrightarrow{\delta} \emptyset; D_{11}^T \xrightarrow{\alpha_T} D_{11}^T + T$ $O \xrightarrow{\delta} \emptyset; D_{11}^O \xrightarrow{\alpha_O} D_{11}^O + O$ | Basal Rate<br>(R14-R16) | $\emptyset \xrightarrow{m_1} N$ $\emptyset \xrightarrow{m_2} T$ $\emptyset \xrightarrow{m_3} O$ |
| Methylation<br>(R17) | $D_{00}^O \xrightarrow{\gamma} D_m$ | Demethylation<br>(R18) | $D_m + NT \xrightarrow{\theta} D_{00}^O + NT$ |

**Figure 4:**
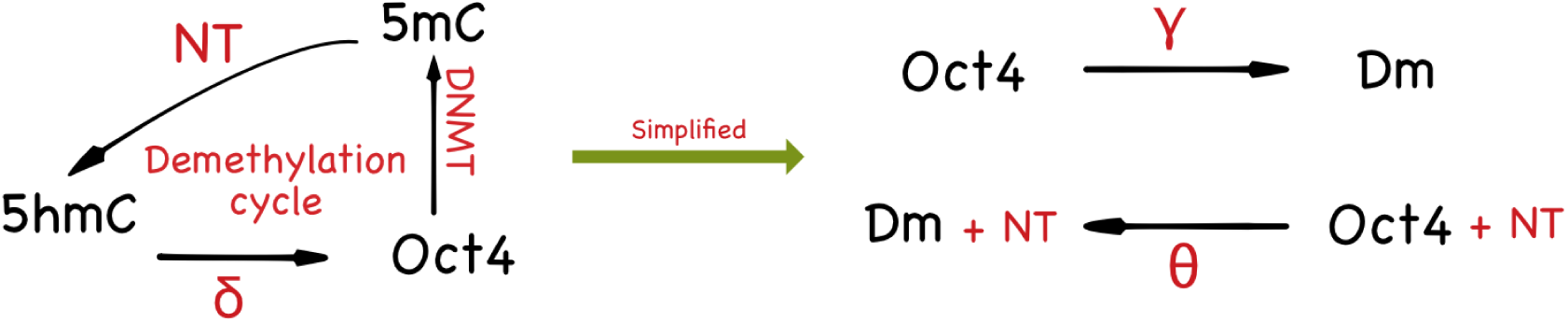
The diagram shows how the triangle topology can be further simplified into a two-species CRN. Since the 5hmC methylation state has a slower dynamics compared to the 5mC methylation state, one may employ a quasi-steady states approximation to reduce the triangle topology to the two-state model depicted on the right.

#### 2.1.1 Full Model Description

##### Description of each module

We grouped our CRN for the full model into eight modules with the reactions numbered from R1 to R18 as shown in Table 1.

1. Oct4 promoter module (R1-R4): This module contains 4 reactions. In our modeling context, we assume that the Oct4 gene has two binding sites. The first binding site is for Oct4 protein itself [52, 24], and the second binding site is for Nanog-TET heterodimer complex [43] as shown in the diamond topology in Figure 3. The parameter *α*_*NT*_ is the forward rate for all reactions that involve NT binding to a promoter, and *K*_*NT*_ is the dissociation rate for all reversible reactions associated with NT binding to a promoter. The parameter *α*_*O*_ is the forward rate for all reactions that involve Oct4 binding to a promoter, and *K*_*O*_ is the dissociation rate for all reversible reactions associated with Oct4 binding to a promoter.
2. TET promoter module (R5-R8): This module contains 4 reactions. Similar to the Oct4 promoter, the TET promoter region is also able to bind with the NT dimer and the Oct4 protein [50, 43, 31]. For the reaction rate, we adopt the same rate parameter for TET as for Oct4.
3. Nanog promoter module (R9): This module describes the reversible reaction of Oct4 protein binding to the Nanog promoter [24].
4. Nanog-TET module (R10): This module describes the fact that the TET-mediated DNA demethylation cycle is guided by the Nanog protein in the form of an NT complex [43]. The parameters *a*_1_, *K*_*d*_ are the association and dissociation rates of the heterodimerization of Nanong and TET.
5. Protein decay and production module (R11-R13): In this module we assume Nanog, TET, Oct4 all have the same degradation rate *δ*. When each gene promoter has been fully occupied by TFs, the corresponding gene will be activated. The protein production rates for each activated promoter 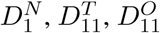 are given by the reaction rates *α*_*N*_, *α*_*T*_ and *α*_*O*_.
6. Protein basal production module (R14-R16): In this module we assume that TET protein has a basal production rate *m*_2_. The rates *m*_1_ (basal rate for Nanog) and *m*_3_ (basal rate for Oct4) are 0. We used the values of such basal protein production rates from [50].
7. Oct4 promoter Methylation module (R17): This module describes the de novo methylation of the Oct4 gene promoter region with a methylation rate *γ*.
8. Nanog-guided TET demethylation module (R18): In this module, we assume that the triangle topology of DNA demethylation cycle has a slow dynamics on the 5hmC oxidized promoter state [31]. Using a quasi-steady state approximation, we reduced the demethylation cycle from the triangle topology to a two-state CRN with the effective demethylation rate given by *θ*. (See Figure 4)

##### Model reduction via quasi-steady steady approximation

The full model CRN is computationally intensive to simulate and conceptually difficult to interpret its dynamical behavior. Biologically speaking, it is often the case that the DNA methylation dynamics operates on a slower time scale than the protein level’s dynamics [44]. Therefore, we make the following reasonable assumptions.

We assume that the reaction rates of the promoter-TF binding and unbinding dynamics are faster than the other reaction rates in the CRN Table 1. By applying time scale separation or quasi-steady state approximation, we reduce the full dynamical system model from a 17 dimensional state space to a 4 dimensional state space. Note that, unlike TF-promoter binding/unbinding, DNA methylation kinetics are as slow as protein kinetics or even slower [44].

The full model ODEs are given in the Methods section. The reduced dynamical system’s ODEs can be written as below:

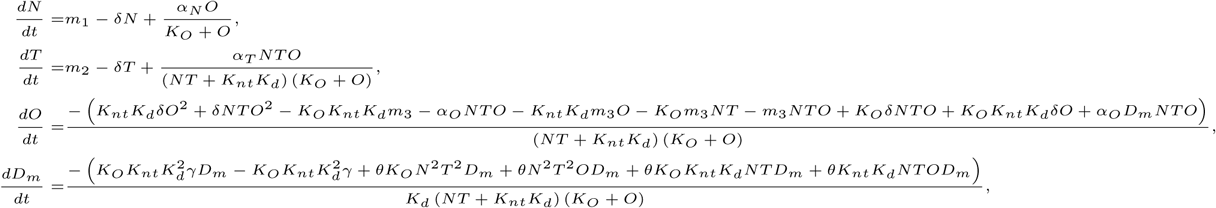

#### 2.1.2 Effect of methylation rate

For the full model, Figure 5 displays how the BoAp changes as one changes the epigenetic reaction rate *γ*. Each plot in the figure corresponds to one set of parameters, sampled from a wide range. The parameter set for the full model contains the following rates: *K*_*O*_, *K*_*nt*_, *K*_*d*_, *a, a*_*nt*_, *a*_*O*_, *α*_*T*_, *α*_*O*_, *α*_*N*_, *δ, γ, θ, m*_1_, *m*_2_ and *m*_3_. Biologically speaking, changing the reaction rate *γ* is equivalent to changing the effective methylation rate. Our subsequent study of the single gene model will display the same trend of BoAp’s dependence on *γ*. The higher the effective methylation rate, the more likely it is for the underlying dynamical system to stay at the somatic cell state, which is represented by a larger BoAp. The phenotypic somatic state is characterized [50] by low expression of Oct4, Nanog, TET, and methylated promoters. With simulations performed over 6200 parameter sets as shown in Figure 5, the BoAp vs. *γ* plot shows that the BoAp is monotonically increasing with respect to *γ*. Note the different behavior for intermediate values of *γ* compared to low or larger values of *γ*. Another interesting observation is that the BoAp is bounded below by 10% for all parameter sets when *γ* is greater than 10^−1.5^.

**Figure 5:**
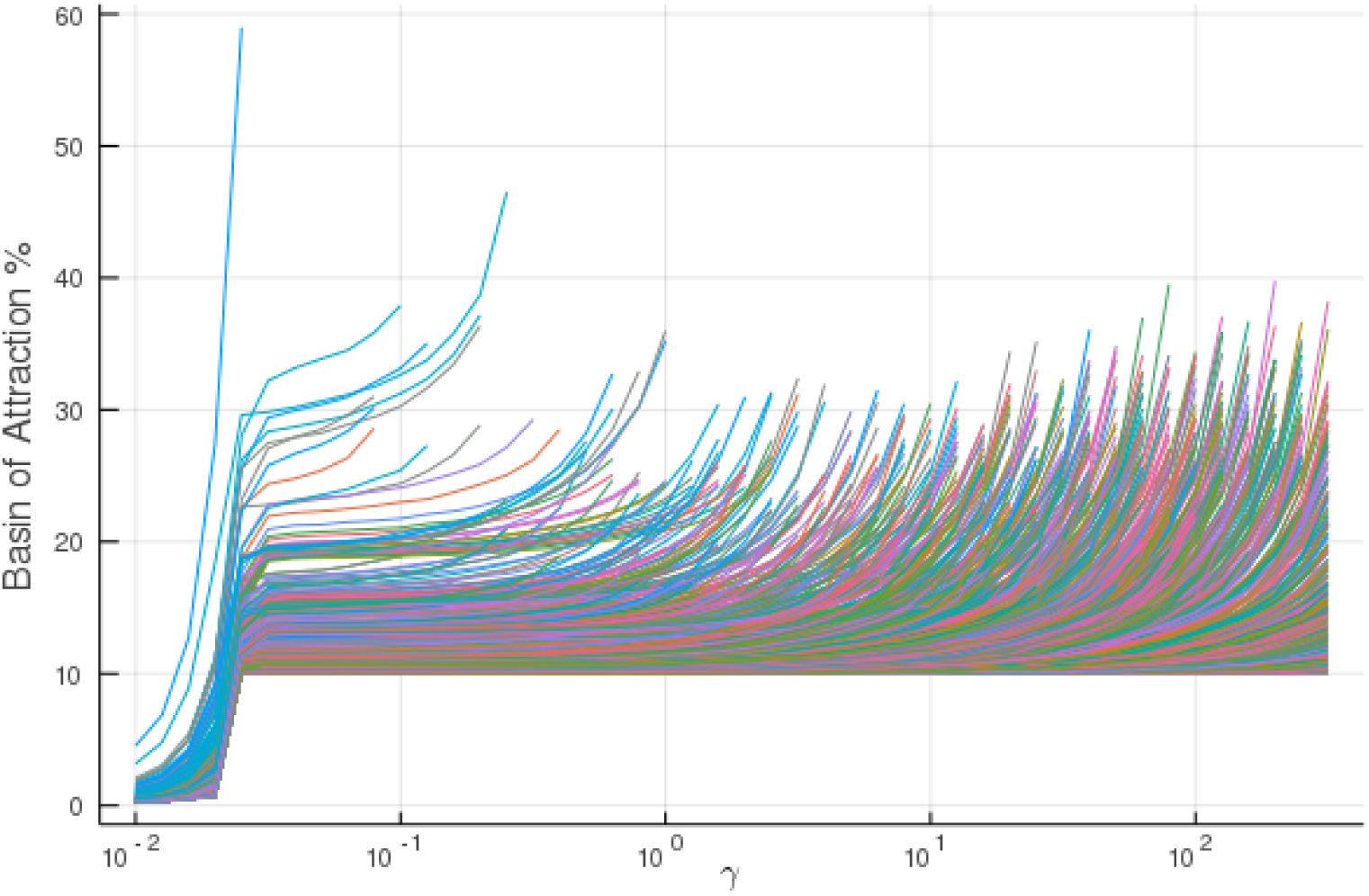
BoAp vs *γ* simulated with a large sets of different rates for the full model. The BoAp increasing rate exhibits a sudden slowing down at around 10^−1.5^

On the other hand, changing the reaction rate *θ* is equivalent to changing the effective demethylation rate. As we reduced the triangle topology for the DNA demethylation cycle to a two-state cycle in the full model, the effective demethylation rate can be thought of as representing the combination effect of 5mC to 5hmC oxidation rate affected by the NT complex and the rate of oxidized 5hmC reverting to the unbound promoter state via either TDG-BER pathway or dilution via cell replication. In Figure 6, the simulation shows that the BoAp is monotonically decreasing with respect to *θ*. Over 6200 parameter sets have been simulated and we plot BoAp vs. *θ* over 4 orders of magnitude to show the trend. We choose the value of the range for *θ* to be the same as the range of *γ* in the BoAp vs. *γ* plot, which both are from 10^−2^ to 10^2^.

**Figure 6:**
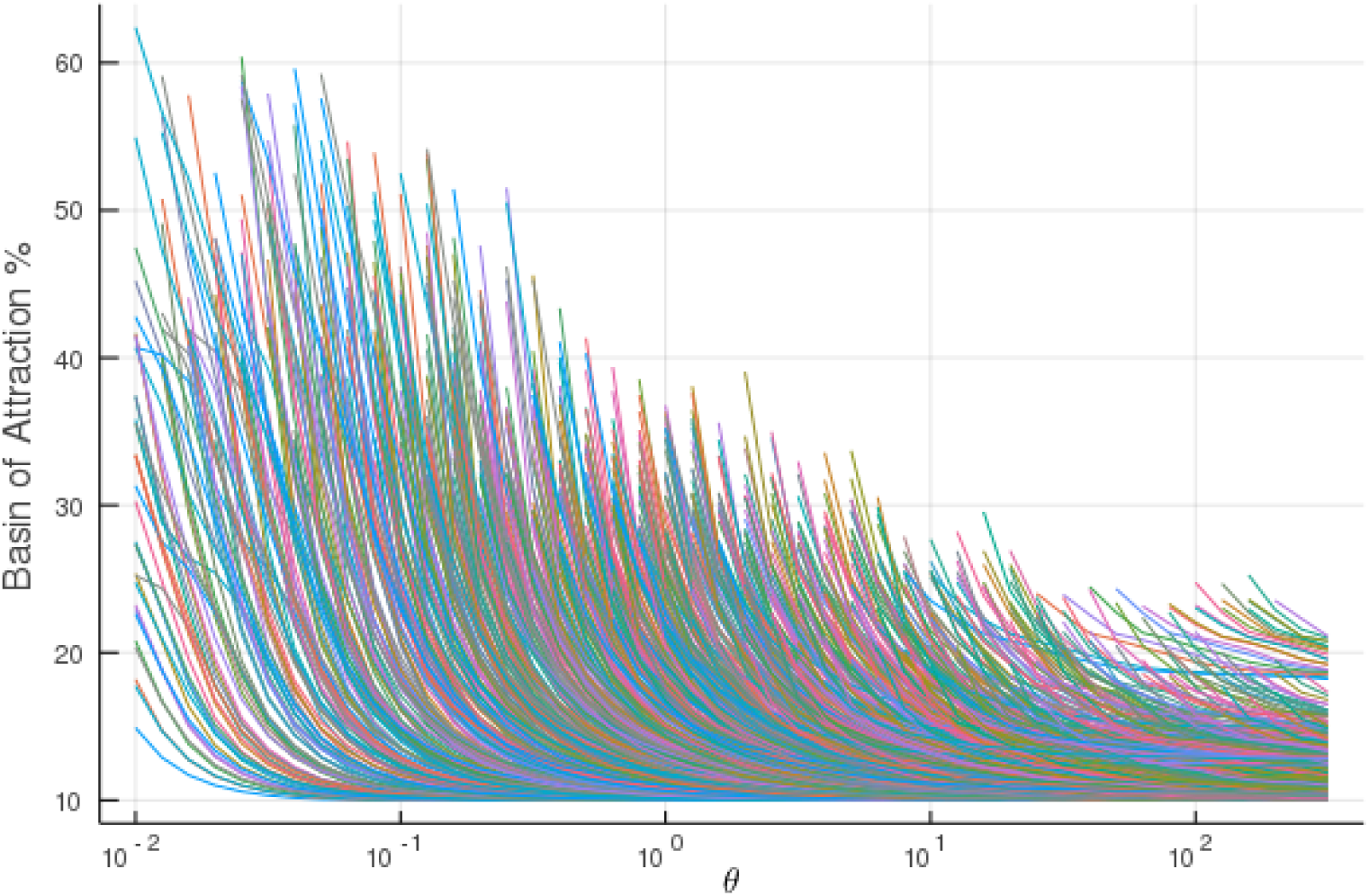
BoAp vs *θ* for a large set of different rates for the full model. This plot shows BoAp is monotonically decreasing with respect to the effective demethylation rate *θ*.

These results confirm that our model captures the key role that methylation plays in enhancing the stability of the somatic state.

#### 2.1.3 Effect of slow/fast methylation kinetics

To understand how the relative methylation rate affects the BoAp, we again define the methylation association rate as 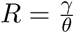. We simulate the full model across different fixed ratios of *R*, letting *γ* range over 4 orders of magnitude.

The simulations in Figure 7 show that BoAp vs. *γ* has a biphasic trend across 5 orders of magnitude in *γ* while *θ* is increased so as to keep the ratio *R* constant with different ratios of *R* that span 3 orders of magnitude.

**Figure 7:**
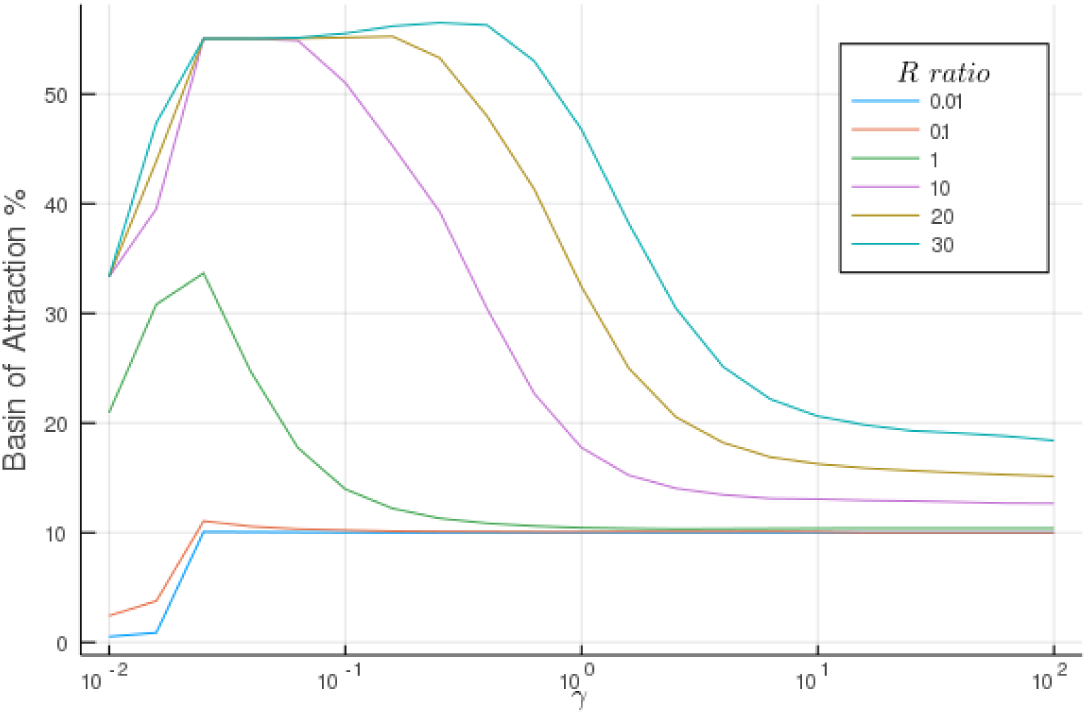
BoAp vs *γ* while *θ* is increased so as to keep the ratio *R* constant. The kinetic rate parameters are: *K*_*O*_ = 0.3, *K*_*nt*_ = 0.2, *K*_*d*_ = 0.1, *a*= 1, *a*_*nt*_ = 100, *a*_*O*_ = 100, *α*_*T*_ = 1, *α*_*O*_ = 1, *α*_*N*_ = 1, *δ* = 1, *R* ∈ {0.01, 0.1, 1, 10, 20, 30}, *γ* ∈ [10^−2^, 10^2^].

The biphasic nature of the trends is not a result of high-dimensional dynamics. Rather, we will reobserve the same behavior in our subsequent study of the single gene model. Furthermore, the biphasic trend for the fixed ratio *R* can roughly be inferred from the PSCC simulations with varying each *γ* and *θ* independently. As per the trend we shown in Figure 5, the BoAp increasing rate exhibits a sudden slowing down at around 10^−1.5^, while there is no corresponding change for the demethylation rate *θ* shown in Figure 6. Therefore, it is not surprising that the methylation association ratio *R* has a biphasic behaviour.

### 2.2 Model of a single self-activating gene

In order to complement the numerical investigations in the previous sections, we consider next a simplified model which includes Oct4 only. Its interaction with the rest of the network is modeled as a self-activating loop. This model is more amenable to an analytical study as will be discussed below.

Our simplified model consists of a single self-activating gene. A CRN model for it can be written as follows:

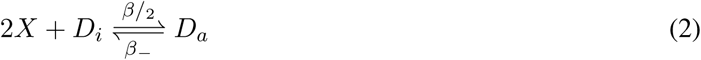

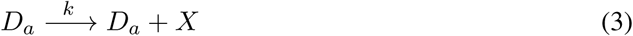

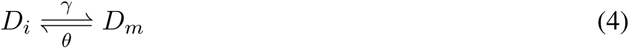

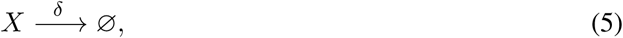

where *X* is the protein expressed from the active promoter state *D*_*a*_ and *D*_*i*_ is the inactive promoter state. (As earlier, we omit explicit variables for mRNA and other intermediate steps such as protein maturation and post-translation modifications.) To model self-activation, we assume that an *X* homodimer can bind to the inactive promoter site to activate the gene. The inactive promoter region can be methylated to a state that we denote by *D*_*m*_. Reversely, *D*_*m*_ can be epigenetically modified back to the *D*_*i*_ state. We assume that protein *X* has degradation rate *δ*. Our aim is to study the effect of the methylation ratio *γ/θ* on the dynamics.

Since the total concentration of promoters is conserved, the ODE model can be reduced by writing *D*_*i*_ = 1 − *D*_*a*_ − *D*_*m*_, where we scale the total number of promoters to 1. The system of ODEs that results is given as follows:

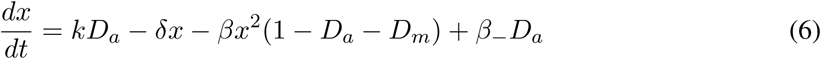

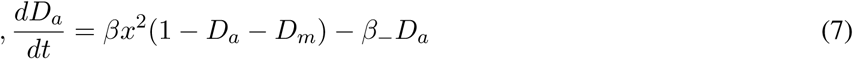

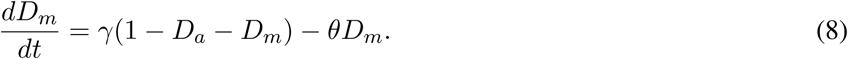

The steady states can be evaluated analytically to yield a cubic equation. There always exists a steady state in which *x*=0 (and also *D*_*a*_=0). We call such a state the *silenced steady state*. It is easy to verify by linearization that this state is locally stable, for any parameter values. A bifurcation occurs and three steady states exist if and only if the following inequality holds:

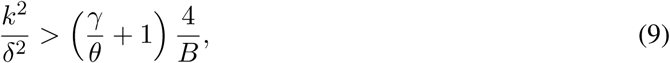

where *B* := *β*_−_*/β* is (double) the dissociation ratio, and *k/δ* and *γ/θ* are the production and methylation ratios respectively. One of two additional steady states is always locally asymptotically stable and the other one is unstable (a saddle). We call the second stable steady state *the active steady state*. It follows from condition (9) that a sufficiently high methylation ratio renders the active steady state non-existent.

The model can be reduced further by utilizing the fact that transcription factor *binding/unbinding is fast* relative to protein expression and epigenetic modifications. We thus carry out a quasi-steady state (QSS) approximation, in which we set the state *D*_*a*_ to:

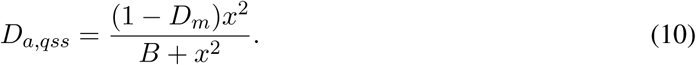

Substituting this expression into the ODE system, we obtain a two-dimensional system whose two variables are the promoter methylated state *y*(*t*) := *D*_*m*_(*t*) and the protein *x*, as follows:

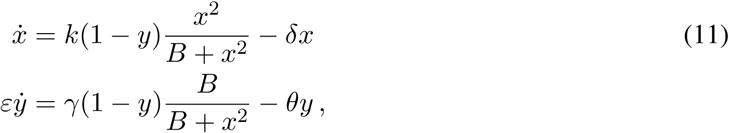

where *ε* is a parameter that represents the relative time-scale separation between protein and methylation kinetics. This system must be studied for (*x, y*) ∈ [0, ∞) × [0, 1]. We next analyze this reduced system.

#### 2.2.1 Global behavior

Observe that all solutions of the reduced system are bounded. Indeed, *y* is bounded by definition, and 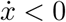 whenever *x* > *k/δ*, so the rectangle [0, *k/δ*] × [0, 1] is an attractor.

The steady states of the reduced system are in a one to one correspondence with the states of the full system, in the sense that for each steady state (*x, y*) there is a unique steady state (*x, D*_*a*_, *D*_*m*_) of the full system, where *D*_*m*_ = *y* and *D*_*a*_ is given by the expression in Eq. (10). Thus there are one or more steady states depending on condition (9). We may think of the system (11) as a “toggle switch” in which *Y* (the methylated promoter) represses *X* (protein), and conversely *X* represses *Y*. If silencing is stronger, we expect the protein state to remain at zero, while if activation is stronger, we expect protein values to converge to a higher steady state.

In order to analyze the global behavior of system (11), we first evaluate its Jacobian matrix:

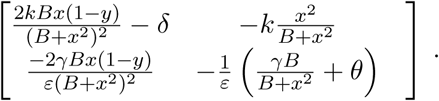

Both off-diagonal entries are non-positive, which makes this system a monotone system, and specifically a system that is cooperative with respect to the cone defined by the (−, −) orthant, as discussed in [45] (see also [46] for an exposition and biological applications). In [26], M.W. Hirsch proved (Theorem 2.2) that every bounded solution of a planar cooperative system converges to an equilibrium; see also Theorem 2.2 in [45]. Coupled from boundedness of solutions for our system, we conclude that every trajectory converges to an equilibrium (which depends on the initial conditions of the trajectory).

##### Phase plane analysis

Our two-dimensional reduced model can be analyzed via nullclines. A summary of the analysis is depicted in Figure 8 for a bistable model. The nullclines and the steady states are independent of *ε*. On the other hand, the separatrix, which is the boundary between the BoA’s, is *ε*-dependent. It is shown that each BoA can be partitioned into *ε*-dependent and *ε*-independent regions. Figure 8-b) shows that the area of the *ε*-independent BoA of *s*_0_ increases as the methylation ratio is increased.

**Figure 8:**
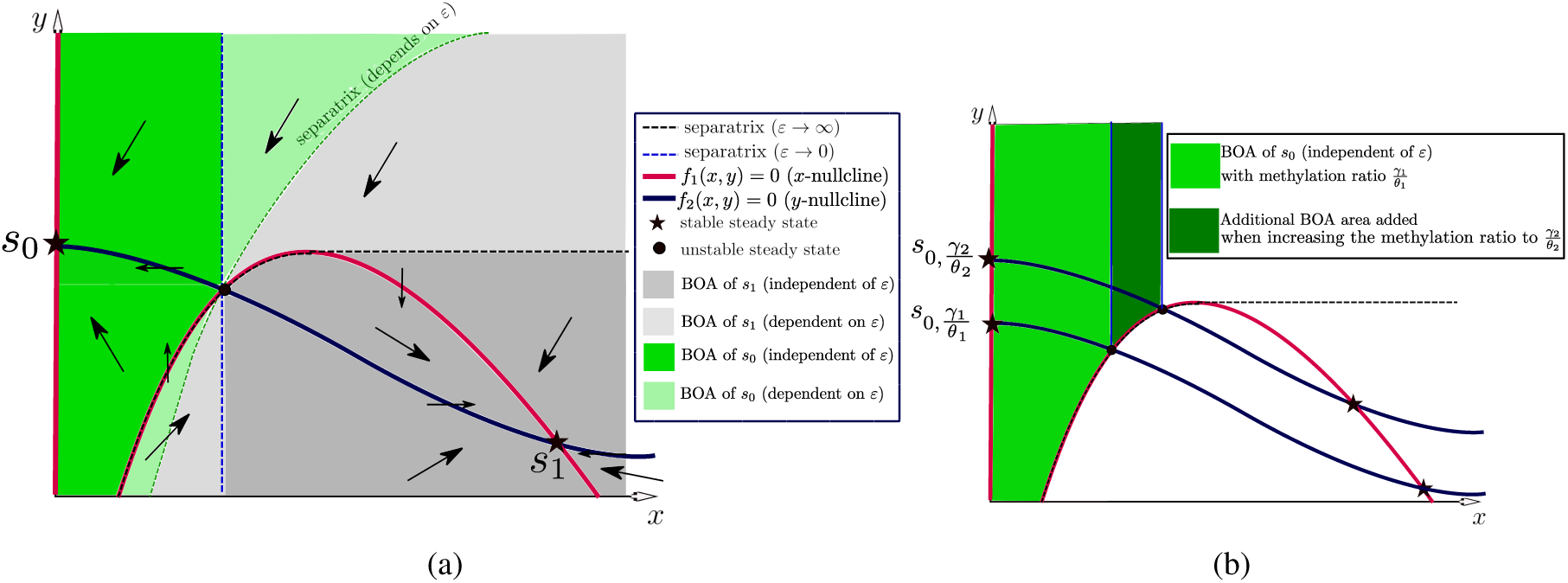
Phase plane analysis of the two-dimensional model (11). (a) The directions of the vector field and the BoAs in the case of bistability with stable steady states *s*_0_, *s*_1_, where *s*_0_ denotes the silenced steady state and *s*_1_ denotes the active steady state. The arrows in each region denote the direction of the vector field. (b) The area of the *ε*-independent BoA of the silenced steady state *s*_0_ increases when the methylation ratio increases.

#### 2.2.2 Slow methylation kinetics

Since DNA methylation is a slow process, we focus on case *ε* → ∞ in this subsection. Figure 8 shows that, in this limit, the BoA of *s*_0_ consists only of the *ε*-independent region.

Next, we analytically investigate the aforementioned limit.

##### Computing the asymptotic separatrix

Setting 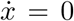, we see that the *x*-nullcline consists of two components, which we call the silenced and active arcs, because the dynamics along each arc converge to the silenced and active steady states, respectively. The arcs can be written as follows:

1. *Silenced:* The arc is *x* = 0, which gives the slow dynamics of

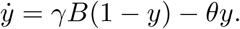 It is a linear system with asymptotically stable steady state at 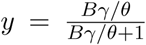. Notice that it is increasing with respect to *γ/θ*.
2. *Active:* The arc is given by the implicit equation: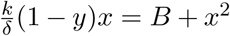. Solving for *x* in terms of *y*:

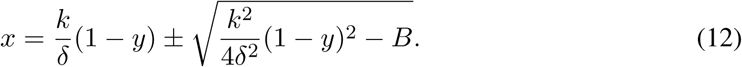

The slow dynamics are given as (note lim sup_*t*→∞_ *y*(*t*) ≠ 1 for all *t* and any initial condition with *y*(0) ≠ 1):

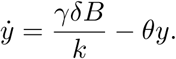

Hence,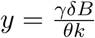 is the asymptotic steady state which is proportional to *γ/θ*.

Given an initial condition (*x*(0), *y*(0)), we want to determine which arc will be representative of the dynamics. A necessary condition for the existence of the second arc is that the quantity under the square root in (12) is real. Hence, we have:

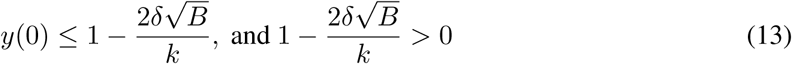

Hence, 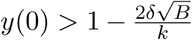 automatically implies that the silenced arc represents the dynamics regardless of the initial *x*(0).

Assuming the necessary condition is satisfied, we need also that

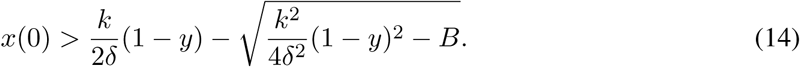

To summarize, if both (13), (14) are satisfied, then the second arc is representative of the dynamics; otherwise, the first arc represents the dynamics. This conclusion is summarized in the dark gray region in Figure 8-a.

##### A remark concerning reprogramming by over-expression of *X*

Our global analysis allows us to intuitively understand the role of *X* in over-expression experiments. In this scenario, we assume that 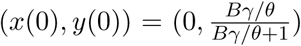, i.e, the initial condition is a silenced steady state. Over-expression means that a sudden dose of *x* will be injected to the system such that *x*(0_+_) = *u*, where *u* is a constant positive number. Our aim is to drive the system to the BoA of the active steady state.

It follows from the analysis above that

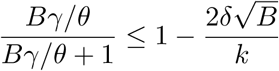

is necessary for reprogrammability. Hence, the expression ratio 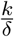 and the methylation ratio 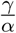 must satisfy:

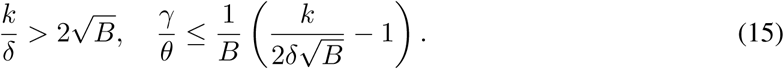

If both of these conditions are satisfied, then overexpression will steer the trajectory to the active arc if the following holds:

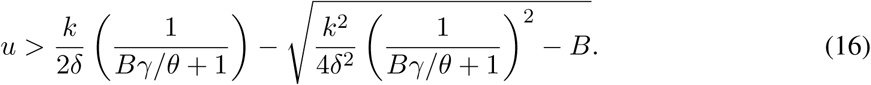

*In a nutshell*, *eqns.* (15), (16) *imply that we need a sufficiently high expression ratio, a sufficiently high overexpression value, and sufficiently low methylation ratio.*

#### 2.2.3 Numerical simulation results for the single-gene model

As a complement to the theoretical discussion, we carried out numerical experimentation to confirm the preceeding nullcline analysis. Figure 9 shows, for the three dimensional system defined by (6) - (8), the result of simulations over 4,000 different parameter sets. The parameter set for the single gene model includes *β, β*_−_, *k,γ, θ* and *δ*. In Figure 9, each single curve corresponds to a fixed parameter set. We observe that, as expected, the BoAp is a monotonically increasing function of *γ*. Intuitively, *γ* is considered to be the effective methylation rate of the Oct4 gene promoter. The methylation rate of the promoter is proportional to the loss of activity of the gene, and hence is proportional to the likelihood of cells staying at their silenced state (defined as the Oct4 protein staying at low concentration level) We view the variable *X* as representing Oct4 in the CRN of the single gene model. We interpret a low concentration of the Oct4 protein as meaning that the system stays at silenced steady state (somatic cell state). On the other hand, if Oct4 protein achieves high concentration, the system is considered as being in the active steady state (the pluripotent state). Conversely, if we increase the reaction rate *θ*, the effective demethylation on the promoter dominates, and therefore a cell would be more likely to stay in a pluripotent state.

**Figure 9:**
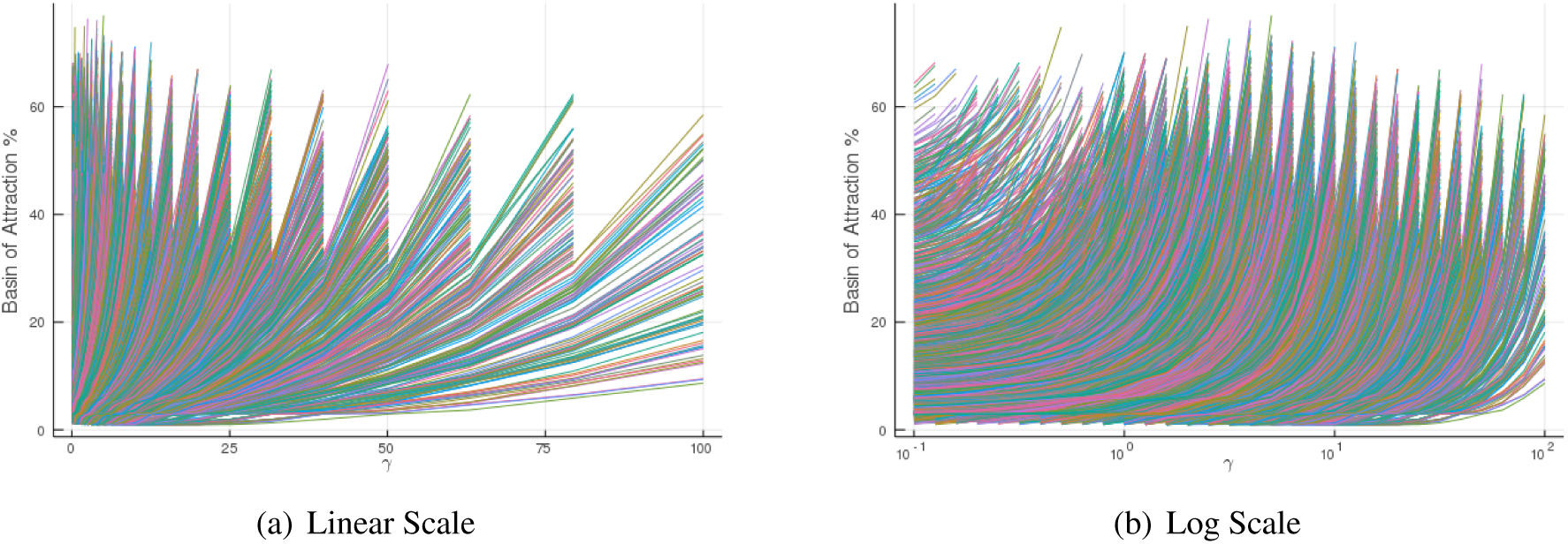
Plots of BoAp vs *γ*, for a large set of different parameters for the single gene model. It can be seen that the BoAp is monotonically increasing with respect to the methylation rate.

We display the simulations both in linear scale, shown in Figure 9(a), and in log scale, shown in Figure 9(b), in order to cover different ranges. The apparent clustering of curves in these plots is due to sampling, as closely related parameters give rise to a similar landscape when changing *γ*. Observe that subsets of curves in the same group all seem to end at roughly the same value of *γ*. The reason for this sudden cut-off is that the the system is undergoing a bifurcation and losing bistability. Furthermore, for each curve in the plot, with *γ* sampled according to a given finite grid, hence the BoAp might achieve a higher value with a finer grid. For example, in Figure 10, the left plot with a coarser grid would indicate that the BoAp reaches only 60%. However, on the right plot, done with a finer grid, the BoAp reaches 72%. We picked a grid size that balances the computational cost involved in the simulation of a large ensemble of parameter sets versus the accuracy of estimation of the BoAp.

**Figure 10:**
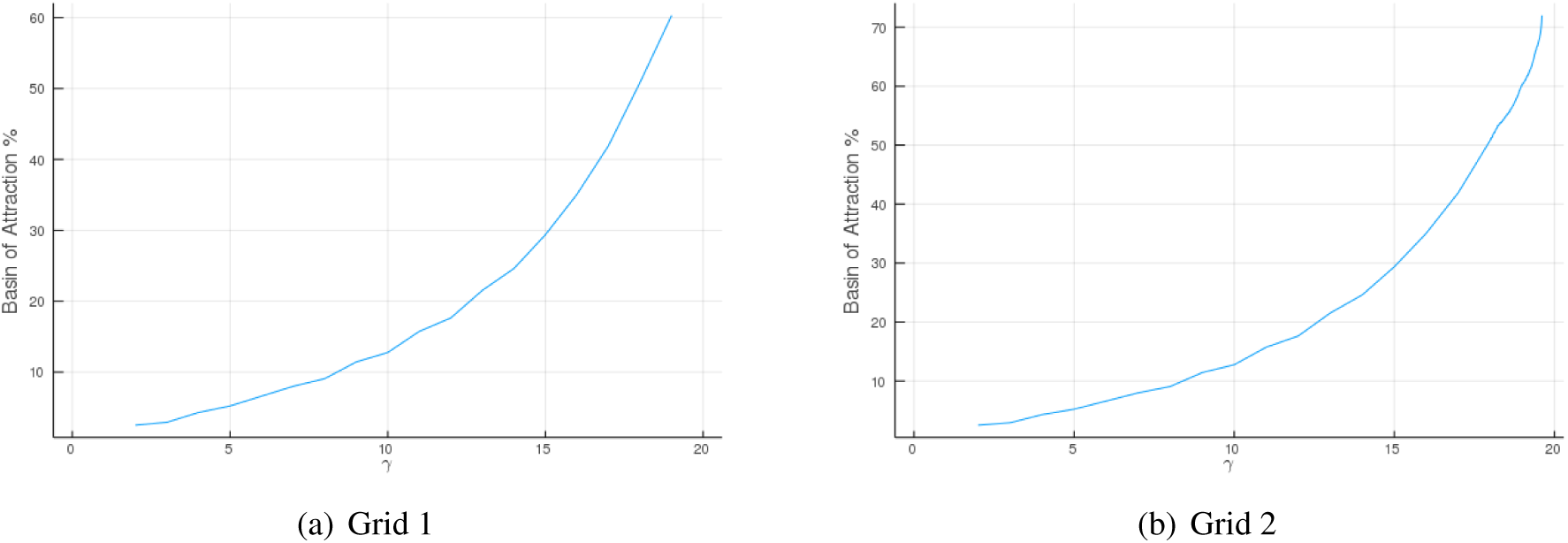
BoAp comparison between different grid sizes. The left plot is simulated with grid size = 1. The right plot has a finer grid size = 0.01. Both plots are simulated with the same parameter set: *β* = 5, *β*_−_ = 5, *k* =5, *δ* = 4, *θ* = 5. We vary *γ* from 0 to 20.

In terms of how the DNA demethylation rate *θ* affects the BoAp, we show a similar simulation in Figure 11 over different parameter sets that for : *β, β*_−_, *k, γ, θ, δ*. We describe the parameter sampling method in the Methods section.

**Figure 11:**
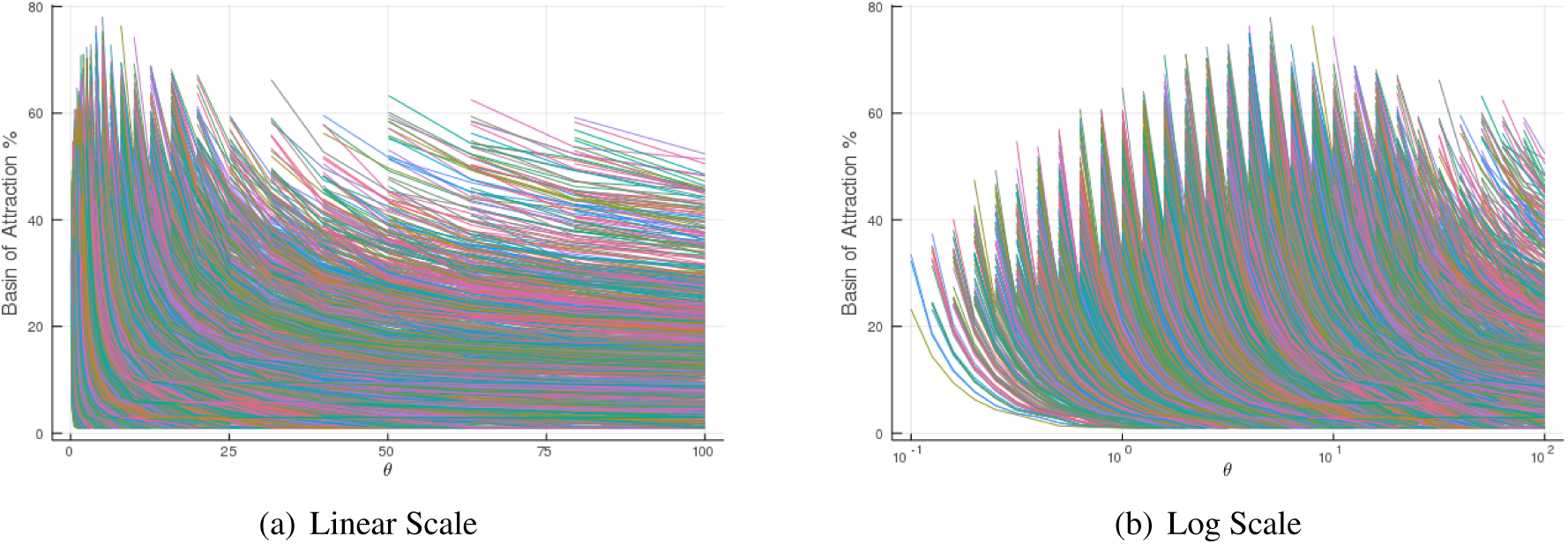
BoAp vs *θ* for a large set of different rates for the single gene model. It can be seen that the BoAp is monotonically decreasing with respect to the demethylation rate.

#### 2.2.4 Effect of slow/fast methylation kinetics

We have discussed in the previous section how the BoAp changes as we independently change either the methylation rate *γ* or the demethylation rate *θ*. Here, we would like understand how the BoAp is affected by the relative changes between *γ* and *θ*. In Figure 12, we explore how the BoAp changes with methylation rate *γ* as one fixes the methylation association ratio defined as

**Figure 12:**
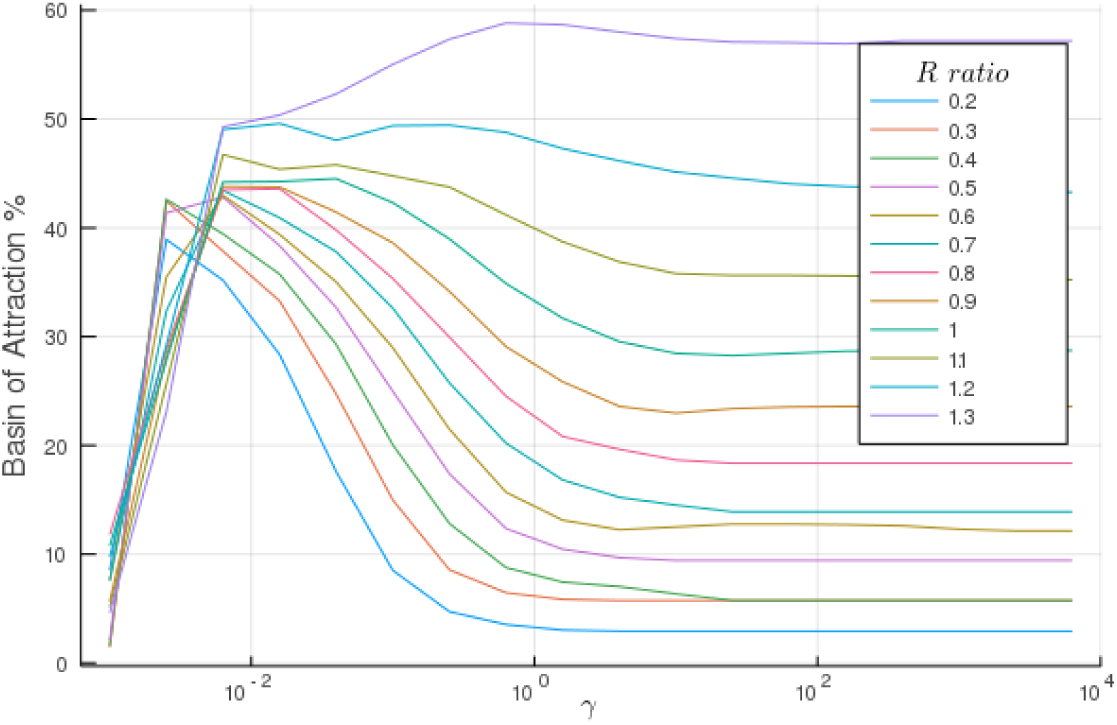
BoAp vs. *γ* with a range of fixed *R* values for the single gene model. For each *R* ratio that allows the bistability of the single gene model, the BOAp vs. *γ* plot has a biphasic nature based on the time scale of the (de)methylation dynamics

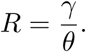

The methylation association ratio (*R*) is the ratio of the methylation to the demethylation rates. The larger the value *R*, the faster the effective methylation process is. Therefore, a value *R* < 1, indicates that the effective DNA demethylation reaction is faster than the methylation reaction. If instead *R* > 1, the effective DNA methylation is a faster process. We explore how BoAp changes as one changes *γ* at different ratios *R*. We study *R* across 12 values and simulate as *γ* and *θ* span across seven orders of magnitudes.

The value of *γ* sets the time scale of the dynamics. When *γ* is small, both methylation and demethylation rate are small. When the value of *γ* is greater than 10^−2^, both methylation and demethylation rate are considered to be large. The simulation in Figure 12 shows a biphasic trend for the underlying dynamics, where *γ* = 10^−2^ set the rough transition barrier. If the (de)methylation rate is sufficiently small, the BoAp is an increasing function with respect to *γ*, and *θ* as well since *R* is fixed. Across different values of *R*, the BoAp shows saturation for large values of *γ*. This saturation trend observed in simulations confirms the theoretical analysis shown in Figure 8. Indeed, as (de)methylation dynamics becomes very fast, the *E* in Figure 8, which controls the time scale of the dynamics, approaches ∞, and the separatrix becomes a vertical line. Therefore, within the predefined cube, the fraction of the volume that corresponds to the somatic state volume will saturate, hence giving rise to the saturation trend in Figure 12.

##### Comparison of the two models

The results for the single gene model parallel those for the four dimensional model (which was, in turn, obtained through QSSA from the full 17-dimensional CRN). For example, we see a qualitatively similar behaviour regarding the BoAp’s dependence on the (de)methylation rate in both models. In addition the results in the reduced model with 4D state space again agree with those for the single gene model with 2D state space.

The full model has the advantage of incorporating all the regulation loops and displaying the effect of the methylation cycle on the steady-state values of the Nanong and TET proteins. However, the single gene model can be reduced to a 2D model which enables the analytical computation of the steady states and the application of phase plane methods. Basic trends in the BoAp can be intuitively read from the shift in the separatrix within the predefined cube. Unfortunately, performing similar analysis for the 4D model is intractable, but the numerical simulations have shown the same basic behavior predicted by the single gene model.

The single gene model is not equipped with the additional regulation loops mediated by the NT heterodimer. Nevertheless, this model demonstrates that the trends displayed by the full model are intrinsic effects of the proposed methylation/demethylation cycle. In both cases, a slow epignetic process –in our case, a meythlation cycle– shifts the stability boundary of the underlying system consistently in the same direction.

## 3 Discussion

We have proposed a GRN model that combines TF-promoter binding/unbinding and gene silencing via DNA methylation. We are currently working on extending the methodology to other types of epigenetic modifications, including histone modifications and chromatin remodeling.

Along with the discovery of the role of TET protein in regulation of the DNA (de)methylation cycle, several theoretical models have been proposed to understand the underlying mechanistic picture of how TFs and DNA interact with each other, and additionally how epigenetic factors such as DNA methylation and histone modifications affect gene regulatory networks in various biological contexts. In the following, we discuss the relation between these models and ours. Many related core gene regulatory networks have been proposed at various levels of complexity. An early attempt using a probabilistic Boolean network that describes the interplay between gene expression, chromatin modifications, and DNA methylation has been proposed in [20]. In addition to lacking mechanistic interpretations, Boolean models do not allow one to quantify the effect of DNA methylation on the stability boundary between the multiple phenotypes. As the role of the TET protein family in gene regulation became clearer [29, 31, 41, 43], various CRN models and ODE models that have partially considered TET-mediated demethylation cycle [50, 1] and core gene-TF regulatory mechanisms have been proposed. Notably in [1], an ODE model for a core gene-TF regulatory network of the PSCC has been studied. The system involves Oct4, Sox2 and Nanog proteins and exhibits multistability under certain conditions. Regarding epigenetic regulation, that work modeled the DNA methylation cycle on top of Oct4 auto-activation. The model did not consider the interaction between Nanog and TET, nor how NT heterodimer regulates the DNA demethylation cycle. Despite the fact that the DNA methylation model in [1] did not include a diamond topology-like mechanism for gene activation, it represented an early attempt to mechanistically link gene regulation and chromatin modifications. In [50], a mechanism was proposed for the core gene-TF network without epigenetics, and then a two-state model without NT regulating the DNA demethylation process was investigated on top of this core network. However, the underlying chemical reaction model in [50] was not spelled-out clearly. In addition to this, without the NT complex involved in the DNA demethylation process, the model in [50] couples the core gene-TF network and DNA demethylation cycle, which does not reflect the experimental evidence of the role of NT in the regulation of DNA demethylation. Specifically, the model lacks the reaction in which NT complex binds to the 5hmC. In the recent experimental work [17], four chromatin regulators, including DNMT3B which causes de novo methylation of cytosine-guanine dinucleotides (CpGs), have been studied. Though a descriptive three-state model was proposed in [17] for explaining how each chromatin regulator affects the silencing and reactivation of gene expression, a detailed mechanism at a molecular level was not provided. In comparison with all these previous attempts, we model the underlying CFN with a detailed CRN model at a molecular level, and we clearly show how epigenetic regulation, specifically the DNA demethylation cycle, regulates the CFN by the Nanog-guided TET protein complex.

In [38], a purely phenomenological model of how epigenetic feedback affects gene expression dynamics is proposed. The main idea is to represent the kinetic rate parameters by epigenetic variables that have their own dynamic equations. More recently, a similar idea has been employed in the context of epithelial-mesenchymal transition (EMT) and cancer metastasis [30]. Such a view of epigenetic feedback regulation of kinetic rates is a valuable complementary perspective.

So far, we have explored a cell fate network combining gene-TF interactions and a TF-guided DNA demethylation process. For the CRN we proposed, we have considered how the TET-mediated demethylation cycle potentially affects the gene regulatory network, and subsequently affects the cell fate decision associated with the transcription factors. We have simplified the demethylation cycle into a two-state cycle. Our mechanistic modeling approach using CRN agrees with the intuition on how DNA methylation affects the cell fate network. The faster the methylation rate the more stable the somatic cell state will be. However, in the detailed molecular picture of the cell fate network that involves Oct4, Nanog, TET and potentially many other transcription factors, there might exist all sorts of self or non-self interactions in the network. For example, if one adds Nanog protein self activation, one would expect such a model to potentially exhibit tristability. Although our model explored the simplest case in which the system gives rise to bistability, the basin of attraction boundary of other models with tristability can still be studied by the relative BoAp within a predefined hypercube.

As for future directions, even though the diamond topology is a mechanism-based model, it does not capture the full complexity of the whole picture of gene activation. The implicit assumption for this activation process is that genes are only activated by TFs binding to promoters. Hence, we are studying the extension of our methods to the analysis of more sophisticated transcriptional control mechanisms, such as the formation of super-enhancers [27]. We are also interested in combining machine learning approaches with our CRN model so as to discover new mechanisms when the relevant data is available. Finally, when generalizing the BoAp to a stochastic setting, the effective landscape of the dynamical system and the mean first passage time between equilibria are the concepts which are most relevant to the analysis of the stochastic stability boundary. Furthermore, stochastic analysis via effective potentials can be used for defining the BoAp and exploring gene-TF interactions with epigenetic regulation.

## 4 Methods

### 4.1 Basic mathematical concepts and definitions

Mathematically, our model is initially described as a Chemical Reaction Network (CRN). CRN’s provide a natural formalism for representing biological interactions, and in particular biochemical processes, and CRN descriptions map into systems of Ordinary Differential Equations (ODEs) using standard procedures.

#### Chemical Reaction Network (CRN) framework

It is essential for our study to keep track of promoter occupancy, since since DNA methylation is commonly understood as a slow process at the level of promoters. Following [2], each gene is associated with promoter states and one or more a protein states^1^. After the model is set up, we later reduce it by the substitution of quasi steady-state approximations of fast variables, based on appropriate relative time-scales.

We briefly review the general formalism, see e.g. [18]. A CRN is specified by a set of species *𝒮* ={*Z*_1_,..*, Z*_*n*_} and a set of reactions *ℛ* = {R_1_, …, R_*ν*_}. A reaction R_j_ can be written as: 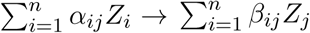. The associated stoichiometry matrix Γ ∈ ℝ^*n×v*^ is defined elementwise as [Γ]_*ij*_ = *β*_*ij*_ −*α*_*ij*_. Each reaction R_j_ can occur with a rate function 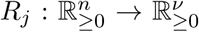. We assume that *R*_*j*_ takes the form of Mass-Action kinetics: 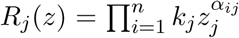 where *k*_*j*_ is a kinetic constant. Letting 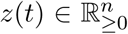 be the vector of species concentrations at time *t*, the associated ODE can be written as: 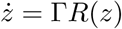, where *R* := [*R*_1_,..*, R*_*ν*_]^*T*^.

The goal of our models is to grasp the essence and the most relevant dynamics that give rise to the CFN with bistability. Unfortunately, it is difficult to decide on appropriate ranges of kinetic parameters for such coarse grained level models. One restriction that we have imposed on parameters is to require bistability, meaning (for both the single gene and the full model) that there should be two stable attractors (and a saddle node) for the underlying dynamics.

### 4.2 Steady State Calculation

For the models defined above (the single gene model is (2)-(5), and the full model given by the CRN in Table 1), we first calculate the number of steady states that the model admits given a certain set of parameters. We use a Homotopy Continuation (HC) method to find all possible steady state solutions for our dynamical system models. The Global HC method is currently the best candidate to find all steady states solution for the system [23].

### 4.3 ODE system for the full CRN model

The corresponding ODE model for the PSCC (Table 1) is given below:

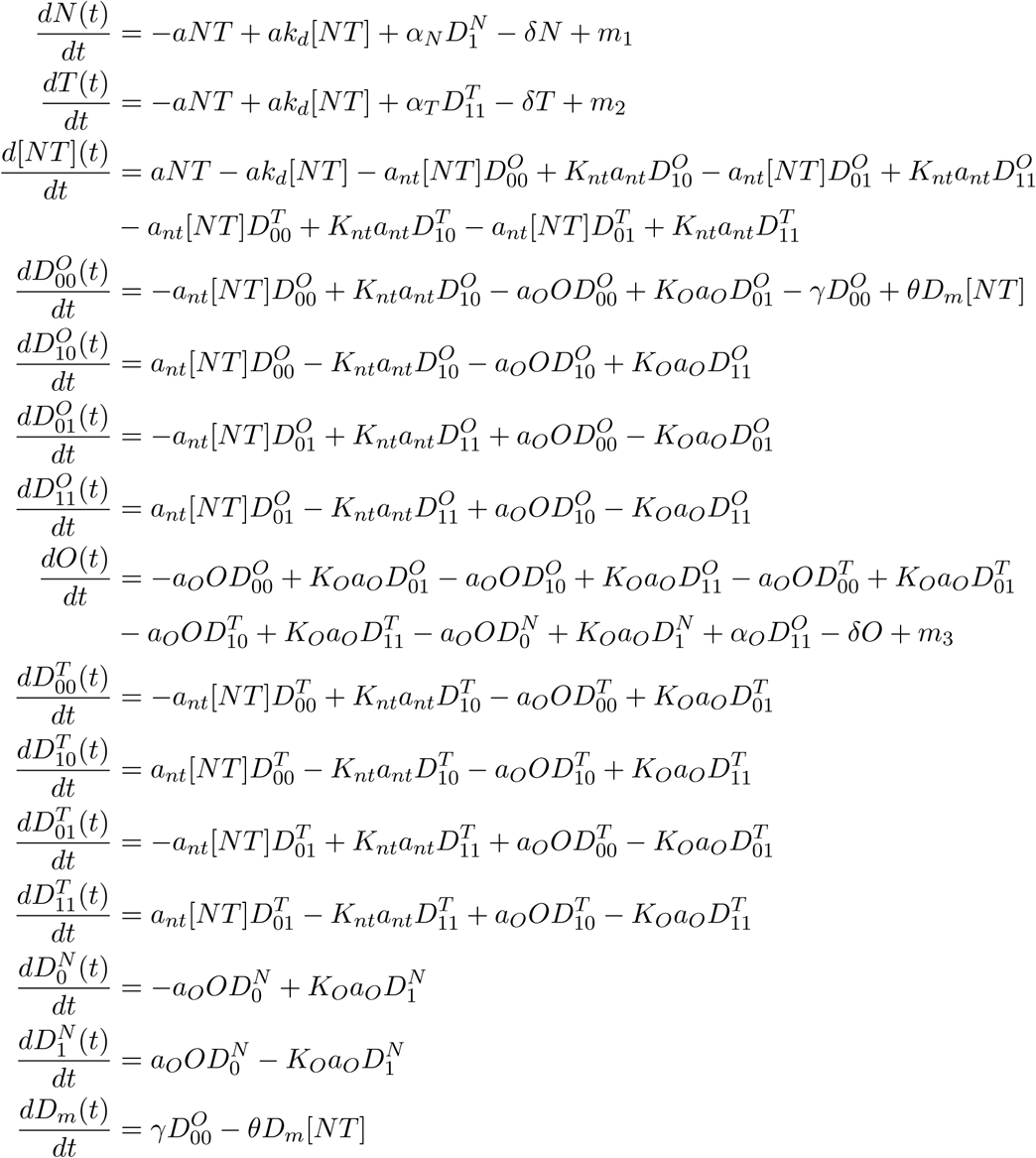

### 4.4 Sampling Method

#### For the single gene model

We calculate the BoAp with various parameter sets that give rise to three steady states. Then, we calculate the BoAp against *γ* and *θ* individually. Though the model contains four state variables *O, D*_*i*_, *D*_*a*_, *D*_*m*_, it is essentially a three-dimensional model due to the conservation of the total concentration of promoters. For simplicity, we normalize total concentration of promoters to an arbitrary constant, say 5.

In order to calculate the BoAp, we need to specify a predefined volume of interest. We have defined the hyper-cube [0, *o*] × [0, *d*_1_] × [0, *d*_2_] where *d*_1_, *d*_2_, *o* are 1.5 times the corresponding high steady state. Because the conservation law puts a constraint on the total concentration of promoters, for each *D*_*i*_, *D*_*a*_ and *D*_*m*_, their concentration should be all less than the total (in the simulation we choose the concentration total = 5). We sample the state space (with a given grid size) and count the number of points in the defined grided space that converges to the low steady state. We then plot BoAp with respect to *γ* and *θ* for various numbers of parameter sets and show that they display a consistent trend.

#### For the full model

Given the CRN from Table 1, we first calculate steady state solutions that show bistability. Then we use quasi-steady steady approximation to reduce the full model from 17 dimensions to 4 dimensions and simulate the BoAp in 4D space with a predefined hypercube similar to the previous paragraph.

### 4.5 Parameters Range

The parameters are constrained to the set which can give rise to bistability.

#### 4.5.1 Kinetic Rates

In the single gene model, the kinetic rate parameters are: *β, β*_−_, *k, δ, γ, θ*. In the simulation of BoAp vs. *γ* plot, we choose the parameter (*β, β*_−_, *k, δ, θ*) values ranging from 0 to 5 with a step size of 0.5. For the simulation of BoAp vs *θ* plot, the same range is used for *β, β*_−_, *k, δ, γ*.

The range can be chosen to be [0, *a*], for any *a* > 0. This is since the ODE (2)-(5) follows Mass-action kinetics and all the rates can be scaled to have values from 0 to *a* for the purpose of BOAp investigation.

In the full model, the kinetic rate parameters that define the system are: *K*_*O*_, *K*_*nt*_, *K*_*d*_, *a, a*_*nt*_, *a*_*O*_, *α*_*T*_, *α*_*O*_, *α*_*N*_, *δ, γ, θ, m*_1_, *m*_2_ and *m*_3_ as shown in the Table 1. Although the values for the three dissociation parameters *K*_*O*_, *K*_*nt*_, *K*_*d*_ were given in [50], in our full model simulation results, we are varying *K*_*O*_, *K*_*nt*_, *K*_*d*_ from 0 to 0.5 with 0.1 step size, and *δ* from 1 to 5 with 0.5 step size, and the parameter *θ* from 1 to 20 with a step size of 2. We set protein production rates *α*_*T*_, *α*_*O*_ and *α*_*N*_ for Nanog, TET and Oct4 to be 1, the parameter *a* = 1, and finally the parameters *a*_*nt*_ and *a*_*O*_ are set to be 100. For those parameters that are set to be constants, we did extensive simulations that have shown that the BoAp is insensitive to large changes in their values. Hence, our presented simulations only contain parameter sets that have noticeable influence on the BoAp.

#### 4.5.2 Methylation association rate (*R*)

In the single gene model, we choose the methylation association rate (*R*) to be from 0.2 to 1.3 so that it covers both the “*R* < 1” regime and the “*R* > 1” regime of the dynamics with a particular parameter set. In the full model, we choose the methylation association rate (*R*) to span across several orders of magnitude, as possible, ranging from 0.01 to 30. The values of *R* for each model are depend on the parameter set we choose, but the qualitative trend for all the parameter sets we tested are the same and show a biphasic trend. In the case of plot in Figure 12, we have chosen the following parameter set: *β* = 4.4, *β*_−_ = 1.8, *k* = 2.5, *δ* = 4.6, and we have varied *R* from 0.2 to 1.3 with *γ* spanning the range from 10^−3^ to 10^4^. For the full model in Figure 7 we have chosen the following parameters: *K*_*O*_ = 0.3, *K*_*nt*_ = 0.2, *K*_*d*_ = 0.1, *a* = 1, *a*_*nt*_ = 100, *a*_*O*_ = 100, *α*_*T*_ = 1, *α*_*O*_ = 1, *α*_*N*_ = 1, *δ* = 1, *R* ∈ {0.01, 0.1, 1, 10, 20, 30}, and with *γ* spanning the range [10^−2^, 10^2^]. For those *R* values outside the range, the system does not exhibit bistability for this particular parameter set.

## 5 Data availability

N/A

## 6 Code availability

The programming code that was used to analyze the raw data that supports the findings of this study is available from the corresponding author upon request.

## Footnotes

1 We assume that transcription is fast enough that no explicit consideration of mRNA abundances is required, but it would be straightforward to add mRNA intermediates to the model.

